# Lsr2, a pleiotropic regulator at the core of the infectious strategy of *Mycobacterium abscessus*

**DOI:** 10.1101/2023.12.12.571305

**Authors:** Elias Gerges, María del Pilar Rodríguez-Ordoñez, Nicolas Durand, Jean-Louis Herrmann, Frédéric Crémazy

## Abstract

*Mycobacterium abscessus* is a non-tuberculous mycobacterium, causing lung infections in cystic fibrosis patients. During pulmonary infection, *M. abscessus* switches from smooth (Mabs-S) to rough (Mabs-R) morphotypes, the latter being hyper-virulent. Previously, we isolated the *lsr2* gene as differentially expressed during S-to-R transition. *lsr2* encodes a pleiotropic transcription factor that falls under the superfamily of nucleoid-associated proteins (NAPs). Here, we used two functional genomics methods, RNA-seq and ChIP-seq, to elucidate the molecular role of Lsr2 in the pathobiology of *M. abscessus*. Transcriptomic analysis shows that Lsr2 differentially regulates gene expression across both morphotypes, most of which are involved in several key cellular processes of *M. abscessus*, including host adaptation and antibiotic resistance. These results were confirmed through RT-qPCR, as well as by minimum inhibitory concentration (MIC) tests and infection tests on macrophages in the presence of antibiotics. ChIP-seq analysis revealed that Lsr2 extensively binds the *M. abscessus* genome at AT-rich sequences and appears to form long domains that participate in the repression of its target genes. Unexpectedly, the genomic distribution of Lsr2 revealed no distinctions between Mabs-S and Mabs-R, implying more intricate mechanisms at play for achieving target selectivity.

**IMPORTANCE:** Lsr2 is a crucial transcription factor and chromosome organizer involved in intracellular growth and virulence in the smooth and rough morphotypes of *Mycobacterium abscessus* (Mabs). Using RNA-seq and ChIP-seq, we investigated the molecular role of Lsr2 in gene expression regulation along with its distribution on Mabs genome. Our study demonstrates the pleiotropic regulatory role of Lsr2, regulating the expression of many genes coordinating essential cellular and molecular processes in both morphotypes. In addition, we have elucidated the role of Lsr2 in antibiotic resistance both *in vitro* and *in vivo*, where *lsr2* mutant strains display heightened sensitivity to antibiotics. Through ChIP-seq, we reported the widespread distribution of Lsr2 on Mabs genome, revealing a direct repressive effect due to its extensive binding on promoters or coding sequences of its targets. This study unveils the significant regulatory role of Lsr2, intricately intertwined with its function in shaping the organization of the Mabs genome.

## INTRODUCTION

*Mycobacterium abscessus* is a rapidly growing mycobacterium (RGM) that has emerged as a significant pathogen in humans (1, 2). This species is known to be one of the most pathogenic non-tuberculous mycobacteria (NTM), representing a serious threat to the health and well-being of cystic fibrosis (CF) patients (3, 4). In comparison to other NTMs, *M. abscessus* infections have been found to have particularly deleterious effects on ventilatory functions (5). It is also responsible for persistent chronic infections associated with the formation of inflammatory granulomas (6). Furthermore, analysis of the *M. abscessus* genome has revealed the presence of several genes associated with multi-resistance to antibiotics and disinfectants which make it hard to treat and eradicate (7–9).

Like other mycobacteria, *M. abscessus* displays two distinct morphotypes phenotypically different in their motility and their ability to grow biofilm: Smooth (Mabs-S) and Rough (Mabs-R) colony morphotypes (10, 11). The loss of surface-associated glycopeptidolipids (GPL) stands as the hallmark of the Mabs-R morphotype, serving as a critical element in its pathogenic strategy (11, 12). Indeed, the transition from Mabs-S to Mabs-R has been found to be correlated with a significant increase in pathogenicity and modulation of the immune system response (13, 14). Additionally, Mabs-R is associated with increased apoptosis and autophagy in cells, leading to cell death and the formation of aggregates or serpentine cords in the extracellular environment, facilitating the multiplication of the bacteria (15, 16). Thus, understanding the mechanisms behind S-to-R transition that only occurs during pulmonary infections is key for unraveling the pathogenicity of *M. abscessus*. Genomic analysis comparing couples of isogenic morphotypes of Mabs-S and Mabs-R identified several insertions/deletions (indels) and single nucleotide polymorphism (SNP) that impair the correct expression of genes involved in the synthesis (*mps1* and *mps2*) and the transport (*mmpL4b*) of GPL (17). Furthermore, we also identified the gene encoding the Leprosy serum reactive clone 2 (Lsr2) protein as exhibiting significantly higher expression level in Mabs-R, thereby indicating its involvement in the heightened pathogenicity of this morphotype.

Originally identified as an immunodominant antigen in *M. leprae* (18), Lsr2 has subsequently been isolated in other mycobacteria (19, 20) and streptomyces (21). This 12.4 kDa small basic protein belongs to the nucleoid-associated proteins (NAPs), a well-known family of chromosomal organizers and global transcription factors (22). Recent studies have highlighted the pivotal role of NAPs as major contributors to bacterial adaptation in response to changes in environmental conditions and stress, particularly during host infections (23). Lsr2 acts as a functional ortholog of the heat-stable nucleoid-structuring protein (H-NS), sharing its ability to preferentially bind AT-rich sequences, possibly forming rigid oligomers and bridging distant DNA fragments across the genome (24). Lsr2 functions as a pleiotropic regulator, influencing the expression of numerous genes involved in virulence and host-induced stress response (25). Chromatin immunoprecipitation-sequencing (ChIP-seq) studies performed in *M. tuberculosis* have revealed that Lsr2 binds to genes associated with early antigenic secretion (ESX) systems, the biosynthesis of phthiocerol dimycocerosates (PDIMs) and phenolic glycolipids (PGL) cell wall lipids, and encoding antigenic proteins of the proline-glutamate (PE) and proline-proline-glutamate (PPE) family (20). Additionally, Lsr2 has been shown as crucial for *M. tuberculosis’* ability to vary oxygen levels and protect bacilli during macrophage infection against reactive oxygen intermediates (ROI) (26, 27). Unlike in *M. tuberculosis*, Lsr2 is not vital for *M. smegmatis* survival but impacts colony morphology, biofilm formation, and adaptation to hypoxia and antibiotics (19, 28).

We recently showed that in *M. abscessus*, Lsr2 is crucial for intracellular growth of both morphotypes within macrophages and amoeba (29). Deletion of *lsr2* increases the susceptibility of the Mabs-R to oxidative species, highlighting the role of Lsr2 in protecting DNA integrity from ROI during macrophage infection. Furthermore, Lsr2 is essential for the virulence of *M. abscessus* in the zebrafish model and persistence in the lungs of infected mice. Here, we used an integrated functional genomic approach to delve into the function of Lsr2 in both S and R morphotypes of *M. abscessus*. Moreover, we investigated its impact on host adaptation and its contribution to antibiotic resistance.

## RESULTS

### Differential expression analysis of the Mabs-S and Mabs-R transcriptomes of *M. abscessus*

We first confirmed our precedent results by performing RNA-seq and comparing gene expression on the S/R pair of 19977-IP (CIP) wild-type control strains. As shown in a prior study conducted by Pawlik and co-workers (17), our data revealed only a handful of genes showing differential expression between the two morphotypes. Specifically, we observed that 4 genes were more expressed in Mabs-S while 9 genes were more expressed in Mabs-R. **(Supplementary Table S2)**. Notably, among these differentially expressed genes, we identified key players in the S/R transition, such as the genes of GPL locus (*mps1, mps2 and gap).* Our current RNA-seq study also unveiled decreased expression of GPL locus genes in Mabs-R compared to Mabs-S. This finding aligns with our previous study, which reported the absence of mRNA transcripts for the *mps1-mps2-gap* operon in the Mabs-R due to the CG insertion in the 5’ region of *mps1*. Furthermore, both our work and the aforementioned study highlight differential regulation of the genes encoding the Nrd family proteins between the two morphotypes. Specifically, transcript expression levels of *nrdF, nrdE, nrdI* and *nrdH* were significantly increased in Mabs-R. These genes are known to be involved in the nucleotide metabolism pathway as well as in the persistence of mycobacteria within phagolysosomes.

### Differential gene expression analysis of wild-type and *lsr2* mutant strains for Mabs-S and Mabs-R

We performed differential gene expression analysis between *lsr2* mutant and wild-type strains to identify the Lsr2 targets in *M. abscessus*. Our RNA-sequencing analysis revealed that a total of 215 genes were regulated by Lsr2 in Mabs-R and 385 genes in Mabs-S. Interestingly, more than 50% of these genes were upregulated in both morphotypes (158 genes in Mabs-R and 220 genes in Mabs-S) in the *lsr2* mutant, providing further evidence for the repressor role of Lsr2 under optimal growth conditions (**Figure 1A**). Among those, only 126 genes were common to both Mabs-S and Mabs-R (**Figure 1B**). These include genes that are either repressed (e.g., *MAB_2037, MAB_2275, MAB_4467*…) or activated in both morphotypes (e.g., *eis2, paaK, MAB_2355c…*) but also genes that show opposite regulation (e.g., *mmpL8, fadD9, MAB_0829…*). We also show that 67.2% and 41.4% of the deregulated genes are only regulated in Mabs-S and Mabs-R respectively (e.g., *mps2, MAB_1012c, ispe…*). This global analysis suggests a different regulatory role for Lsr2 in the two morphotypes depending on their physiologies.

**Figure 1.**
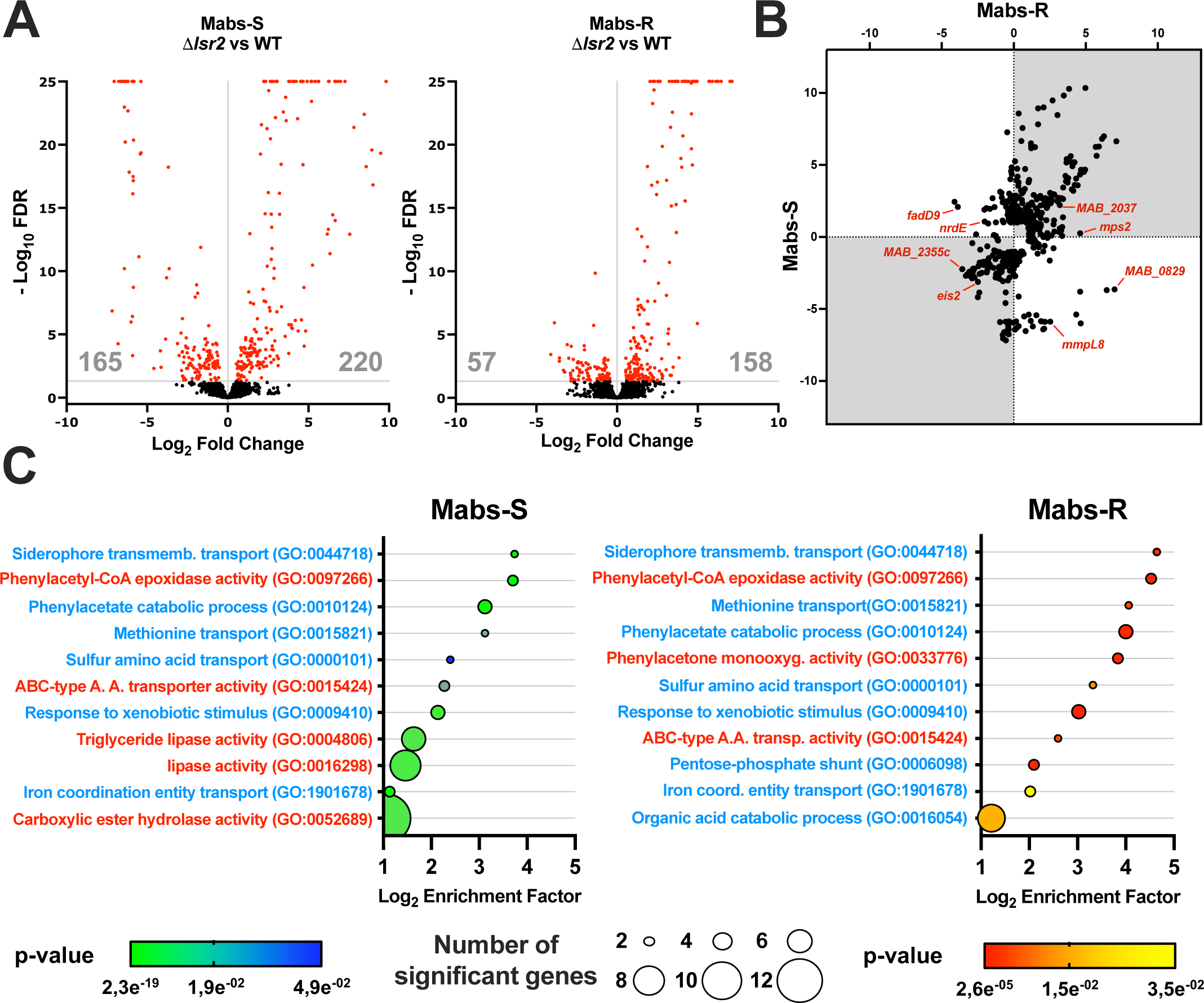
Analysis of Lsr2 regulon in *M. abscessus* morphotypes. **A.** Volcano plots showing differentially expressed genes (DEG) (red) in Mabs-S-Δ*lsr2* vs Mabs-S-WT (left) and Mabs-R-Δ*lsr2* vs Mabs-R-WT (right). Genes exhibiting a greater than 2-FoldChange in expression between two conditions (Log_2_ FoldChange ≤-1 or ≥1) and with a false discovery rate (FDR) of less than 0.05 are considered significantly deregulated. Genes with Log_2_FC ≥1 correspond to down-regulated genes by Lsr2 (repressed genes) and genes with Log_2_FC ≤-1 correspond to up-regulated genes by Lsr2 (activated genes). **B.** Plot representation showing all genes regulated by Lsr2 at least in one of the two morphotypes of *M. abscessus* with a Log_2_FC ≥1 and Log_2_FC ≤-1 of gene expression between the Δ*lsr2* mutant strains and wild-type strains. **C.** GO enrichment analysis was performed with the topGO R package. Enriched GOs (Biological Process in blue and Molecular Function in red) are sorted according to their enrichment factor (EF), corresponding to the ratio of significant DEGs assigned to the GO over expected assigned DEGs to the GO as defined by topGO. Enriched GOs are represented by circles whose size is proportional to the amount of significant DEGs assigned. Positive statistical tests are given that face each GO.

Gene Ontology (GO) analysis performed on both Mabs-S and R showed that Lsr2 affects multiple categories of biological and functional processes in *M. abscessus*, confirming its role as a pleiotropic transcription regulator (**Figure 1C**). Notably, Lsr2 exhibits a significant impact on the transport of different substrates in both morphotypes including siderophore transport (GO:0044718), methionine transport (GO:0015821), sulfur amino acid transport (GO:0000101), iron coordination entity transport (GO:1901678) and ABC−type amino acid transporter activity (GO:0015424). Pertaining to metabolism activity, we observed GO enrichment associated with phenylacetate catabolism (GO:0097266, GO:0010124, GO:0033776), metabolism of glucose (GO:0006098) and organic acid catabolism (GO:0016054). Interestingly, we observed an enrichment in lipid metabolism (GO:0016298, GO:0004806) exclusively in Mabs-S suggesting a regulatory role for this protein based on the membrane lipid composition of the two morphotypes. In addition, GO related to bacterial stress response is also enriched in both Mabs-S and Mabs-R, particularly in the context of the xenobiotic stimulus (GO:0009410).

### Lsr2 modulates the expression of genes involved in intracellular growth and in response to stress

Our transcriptomic analysis shows that Lsr2 acts as a pleiotropic transcription factor in both morphotypes of *M. abscessus,* regulating a wide range of genes involved in key physiological and cellular processes. Lsr2 positively controls the expression of genes related to growth and intracellular survival (**Figure 2A**), such as *paaA, paaB, paaI, paaJ and paak* genes which were acquired through horizontal transfer from *β-proteobacteria* and involved in phenylacetic acid (PAA) degradation (7). *nrdI* and *nrdE* genes, two key players of intracellular survival (17) are both upregulated by Lsr2 in Mabs-R, while they are downregulated in Mabs-S. Deletion of *lsr2* leads to a significant decrease the expression of *MAB_4532c* (*eis2*) that encodes the Eis N-acetyl transferase protein (30) both in Mabs-S and Mabs-R (**Figure 2A**).

**Figure 2.**
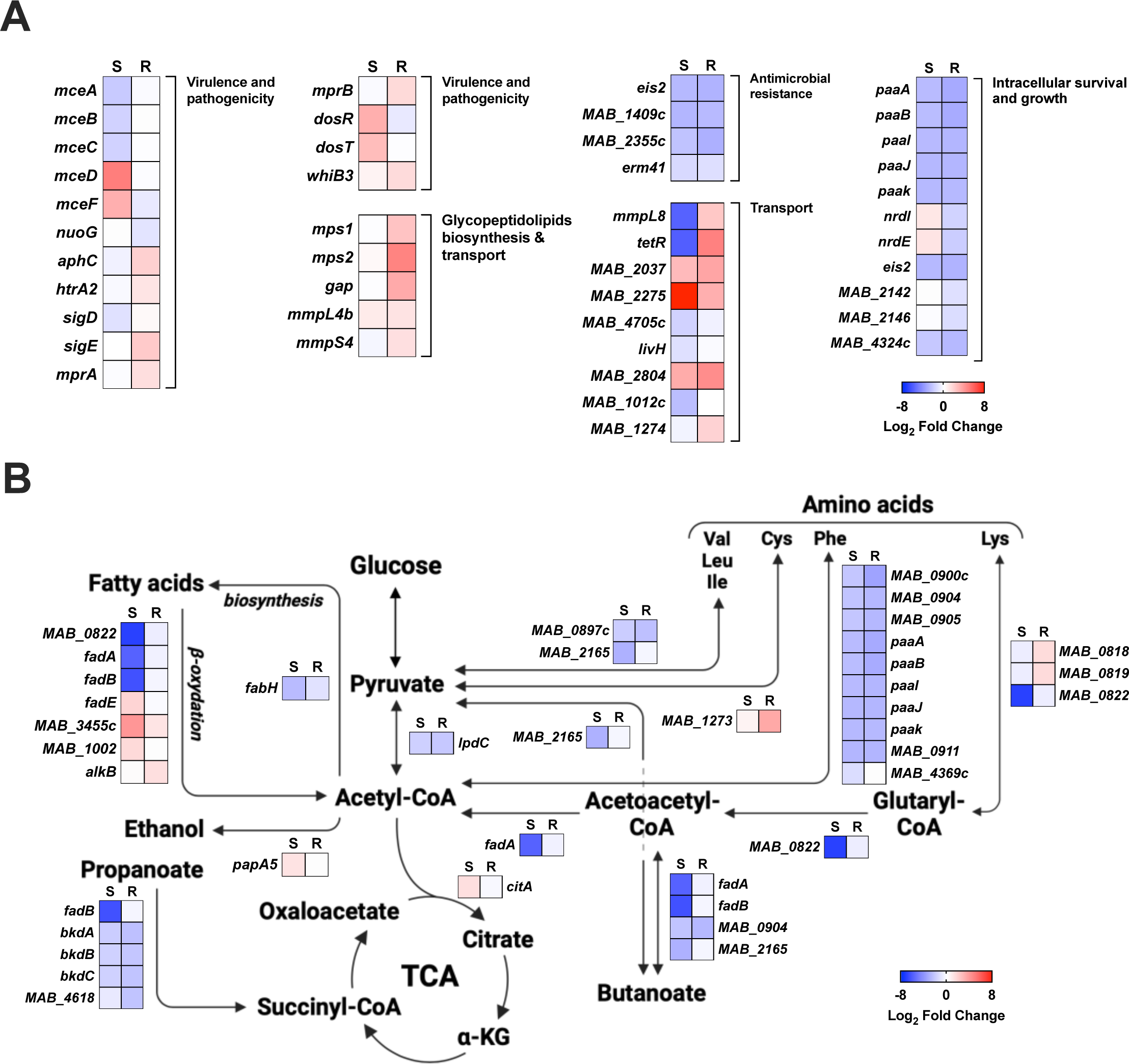
Lsr2 regulated genes involved in different pathways in *M. abscessus*. **A** and **B**. Heat maps representing genes regulated by Lsr2 and implicated in various cellular mechanisms (intracellular survival and growth, virulence and pathogenicity, glycopetidolipids biosynthesis, transport, antimicrobial resistance genes and central carbon metabolism) in Mabs-S and Mabs-R. The color scale represents differentially expressed genes (DEGs) with a Log_2_FC ranging from −8 and 8. Genes with Log_2_FC ≥1 correspond to down-regulated genes by Lsr2 (repressed genes) while genes with Log_2_FC ≤-1 correspond to up-regulated genes by Lsr2 (activated genes).

Additionally, we identified two genes, *MAB_2142* and *MAB_2146* activated only in Mabs-R by Lsr2 and encoding [4F-4S] Ferredoxin proteins that function as intracellular stress sensors and contribute to genomic stability during macrophage and amoeba infections (30). Deletion of *lsr2* in Mabs-S and Mabs-R also results in a significant decrease in the expression of the *MAB_4324c* gene, which encodes an N-acetyltransferase recently found to enhance internalization and intracellular growth of mycobacteria in human macrophages (31) (**Figure 2A**).

### Lsr2 plays a role in virulence and pathogenicity regulation genes

Lsr2 modulates the expression of *mce* (mammalian cell entry) genes encoding proteins involved in virulence and pathogenicity through host cell recognition and invasion in *M.tuberculosis* (32, 33). For example, *mceA, mceB* and *mceC* genes were more expressed in wild-type Mabs-S compared to Mabs-S-Δ*lsr2* while other *mce* genes such *mceD* and *mceF,* located in a different locus, were repressed (**Figure 2A**). *nuoG, aphC* and *htrA2*, all regulated in Mabs-R, impede macrophage antimicrobial effectors through their response to oxidative and nitrosative stress as well as their inhibitory effect on cell apoptosis and phagosomal maturation (34, 35). Interestingly, several families of transcriptional regulators playing a key role in virulence were targets for Lsr2. For example, *sigD* and *SigE* that code for two sigma factors regulated by Lsr2 in Mabs-S and Mabs-R respectively, are involved in macrophage viability through resistance to oxidative stress (36, 37). Moreover, *mprA* and *mprB,* also regulated by Lsr2 in Mabs-R, are *sigD-*dependent transcriptional regulators involved in stress response pathways in *M. tuberculosis* (38). *dosR* and *dosT,* encoding essential factors for hypoxic adaptation, persistence, virulence for *M. tuberculosis*, are tightly controlled by Lsr2 in Mabs-S (39, 40). Finally, Lsr2 controls *whiB3* transcription in Mabs-R, which is known to be involved in intraphagosomal stress resistance in *M. tuberculosis* (41, 42) and redox sensing in *M. smegmatis* (43). The transcriptomic results reveal a diverse range of Lsr2 targets involved in virulence, strongly suggesting its role in the infection, pathogenicity, and persistence of *M. abscessus*.

### Lsr2 regulates families of proteins involved in transport in *M. abscessus*

MmpL proteins (Mycobacterial membrane protein Large) play a crucial role in mycobacterial physiology by facilitating the export of lipid components, toxins, and metabolites across the cell envelope, as well as expelling antibiotics (44, 45). Interestingly, Lsr2 exhibits distinct regulation of the *MAB_0855* gene encoding the MmpL8 protein in both Mabs-S and Mabs-R (**Figure 2A and Supplementary Table S3)**. MmpL8 was found to be essential for intracellular growth of *M. abscessus* in amoeba and macrophages. Its deletion significantly reduces virulence in zebrafish and disrupts interactions with macrophages, hampering the establishment of cytosolic connections (46). In Mabs-S, Lsr2 upregulated the *mmpL8* gene by around 56-fold compared to Mabs-S-Δ*lsr2*, while in Mabs-R, Lsr2 downregulated the *mmpL8* gene by approximately 5-fold in Mabs-R-Δ*lsr2.* To confirm if whether Lsr2 exhibits the same regulatory effect in *vivo*, we performed RT-qPCR analysis on *mmpL8* using mRNA isolated from wild-type and *lsr2* mutants Mabs-S and Mabs-R morphotypes 16 hours after murine J774.2 macrophage infections. Consistent with transcriptomic analysis in planktonic cultures, Lsr2 regulation of the *mmpL8* gene persists during macrophage infections. We observed a significant increase in *mmpL8* transcripts in Mabs-R-Δ*lsr2*, while the expression of this gene was not detectable in Mabs-S-Δ*lsr2* (**Figure 3**). This can also suggest that Lsr2 controls the expression of the *tetR* transcription factor, which in turn regulates the entire *mmpL8_MAB_* locus (**Figure 2A and Supplementary Table S3)**. Additional examples for strongly regulated *mmpL* genes in Mabs-S and Mabs-R were *MAB_ 2037* and *MAB_2075*. The *MAB_2037* gene, belonging to mycolate synthesis operon (30), was significantly repressed by Lsr2 both in Mabs-S and Mabs-R (**Figure 2A**). Again, the repressive effect of Lsr2 on the *MAB_2037* gene was confirmed by RT-qPCR after infection by both morphotypes of *M. abscessus* (**Figure 3**). For all these genes, complementing the mutant strains with *lsr2* restored the same level of expression as observed in wild-type strains.

**Figure 3.**
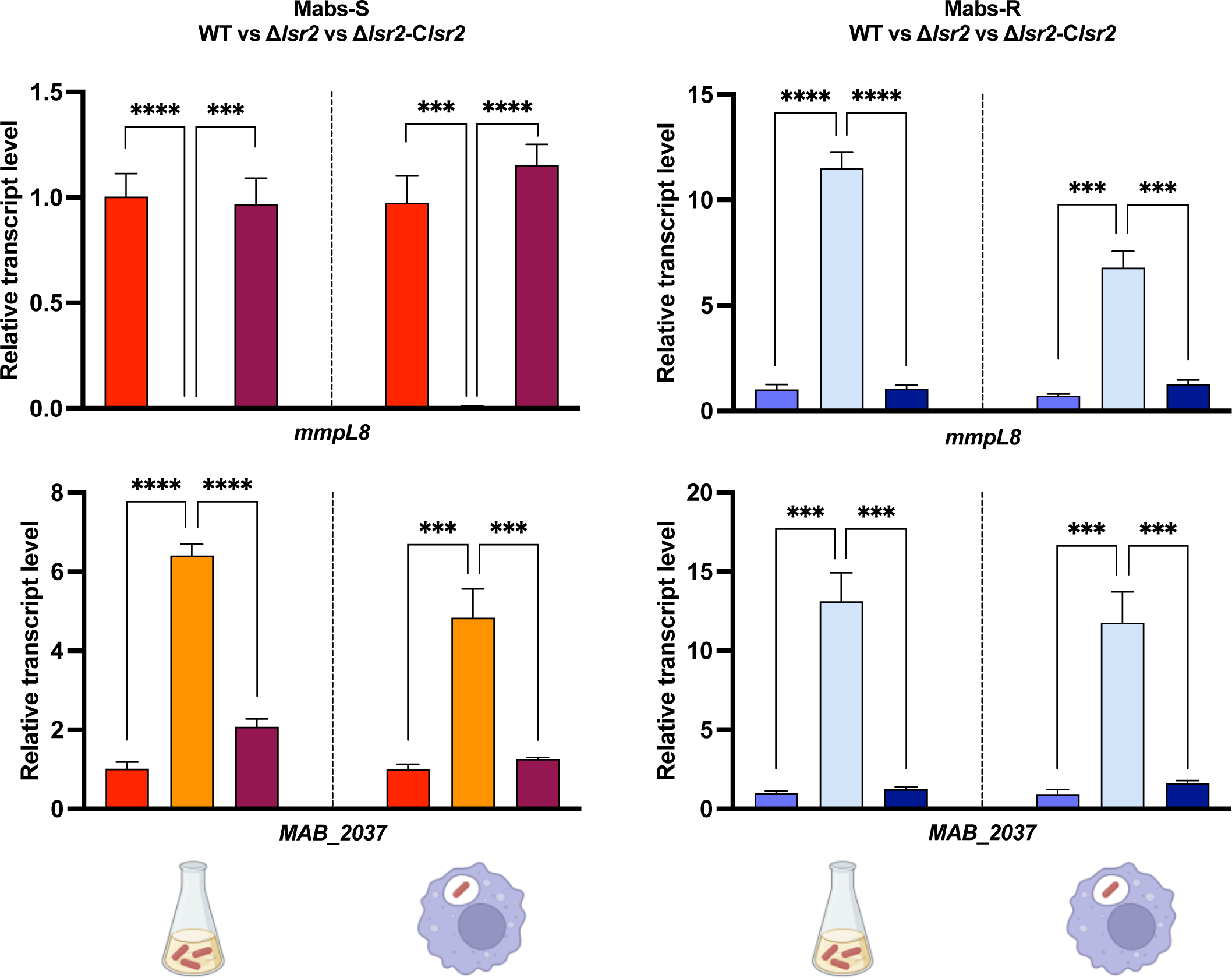
Conservation of Lsr2 regulatory effect on transport genes in both planktonic growth and intracellular growth conditions. Quantification of relative expression of *mmpL8* and *MAB_2037* measured by RT-qPCR under planktonic growth conditions (left side) and intracellular growth conditions (right side) in Mabs-S-WT (red), Mabs-S-Δ*lsr2* (orange), Mabs-S-Δ*lsr2*-C*lsr2* (brown), Mabs-R-WT (blue), Mabs-R-Δ*lsr2* (light blue) and Mabs-R-Δ*lsr2*-C*lsr2* (dark blue). *SigA* transcript levels were used for normalization. For each assay, n= 3 and error bars are SEM. *∗∗∗P < 0.001,* and *∗∗∗∗P < 0.0001* (Student’s *t* test)

### *lsr2* deletion results in increased GPL locus transcripts in Mabs-R morphotype

The loss of GPL at the plasma membrane of *M. abscessus* is known as the hallmark of the S- to-R transition and the role of Lsr2 as a negative regulator of GPL production was already described in *M. smegmatis* (47). However, our previous results showed no effect of Lsr2 deletion on either the GPL profile or the morphology of the bacterium (29). Upon deletion of *lsr2*, genes involved in GPL biosynthesis and transport (*mps1*, *mps2*, *gap* and *mmpL4b*) showed increased expression only in Mabs-R, coinciding with the restoration of a full-length transcript, despite the presence of the genetic lesion, while no significant difference in Mabs-S was observed (**Figure 2A**).

### Lsr2 is implicated in central carbon metabolism regulation

Through KEGG analysis, we investigated the impact of Lsr2 on various metabolic pathways in *M. abscessus* (**Figure 2B**). Notably, our findings revealed that Lsr2 predominantly affects the pathways of fatty acid degradation (ß-oxidation) and amino acid degradation in both morphotypes. Lsr2 exhibits both upregulation and downregulation of numerous genes encoding enzymes involved in ß-oxidation, including *fad* genes encoding acyl-CoA synthetases, ultimately leading to acetyl-CoA production for further incorporation into the tricarboxylic acid (TCA) cycle (**Figure 2B**). The regulatory influence of Lsr2 extends to amino acid metabolism, particularly affecting the *paa* genes involved in intracellular growth, which exhibit increased expression in both morphotypes. Genes involved in propanoate metabolism were mostly highly expressed following *lsr2* deletion in Mabs-S and Mabs-R, including *bkdA*, *bkdB*, *bkdC* and *MAB_4618* that facilitate the conversion of propionyl-CoA derived by β-oxidation of odd-chain fatty acids, into succinate.

### Lsr2 controls antimicrobials resistance genes transcription and sensibility to antibiotics in *M. abscessus*

Among the genes regulated by Lsr2 in both Mabs-S and Mabs-R, we have identified four genes (*eis2, MAB_1409c, MAB_2355c* and *erm41*) encoding proteins modifying the activity of aminoglycosides and macrolides antibiotics families, both widely used in the clinical treatment of *M. abscessus* (9, 48–53). Our RNA-seq data shows that Lsr2 activates these antimicrobial resistance genes, exhibiting higher expression in wild-type strains (**Figure 2A**). Furthermore, RT-qPCR showed a significant decrease of expression of these four genes in Mabs-S-Δ*lsr2* and Mabs-R-Δ*lsr2* upon macrophage infections (**Figure 4A**). Again, the wild-type expression level of these genes was restored by complementing the mutant strains with *lsr2*. To investigate whether this reduction in gene expression correlates with an increase in antibiotics sensitivity, we determined the minimum inhibitory concentrations (MICs) of amikacin, tobramycin and clarithromycin for both wild-type and mutant strains (**Table 1**). Mabs-S-Δ*lsr2* and Mabs-R-Δ*lsr2* were more sensitive to one dilution factor for amikacin (4.5 and 8 µg/mL) and for tobramycin (4.5 and 5.5 µg/mL) compared to wild-type strains. For clarithromycin, Mabs-R-Δ*lsr2* (0.125 µg/mL) was only four times more sensitive than Mabs-R (0.5 µg/mL). Similar experiments performed after macrophage infections followed by treatments with 1xCMI liposomal amikacin and 1xCMI clarithromycin showed that both Mabs-S and Mabs-R respond more significantly to liposomal amikacin and clarithromycin treatments compared to the wild-type strains. For liposomal amikacin, this increased sensitivity is already significant on day 1 post-treatment for Mabs-S (*p < 0.05*) and day 3 for the Mabs-R (*p < 0.01*). By day 6, both mutant strains exhibit a substantial difference in response to liposomal amikacin (*p < 0.0001*). Regarding clarithromycin, the impact of *lsr2* deletion on antibiotic response is particularly prominent in the Mabs-R-Δ*lsr2,* with a significant effect observed as early as day 1 post treatment (*p <0.05*) and a stronger response by day 6 (*p < 0.0001*) (**Figure 4B**). These results strongly indicate that Lsr2, through its transcriptional effect, contributes in lowering susceptibility to antibiotics, thereby confirming its role in antibiotic resistance. Moreover, when assessing intracellular CFU counting following infection without antibiotic treatments, we observed a more important reduction in bacterial viability in the presence of antibiotics for *lsr2* mutant strains (**Supplementary Figure S1)**.

**Figure 4.**
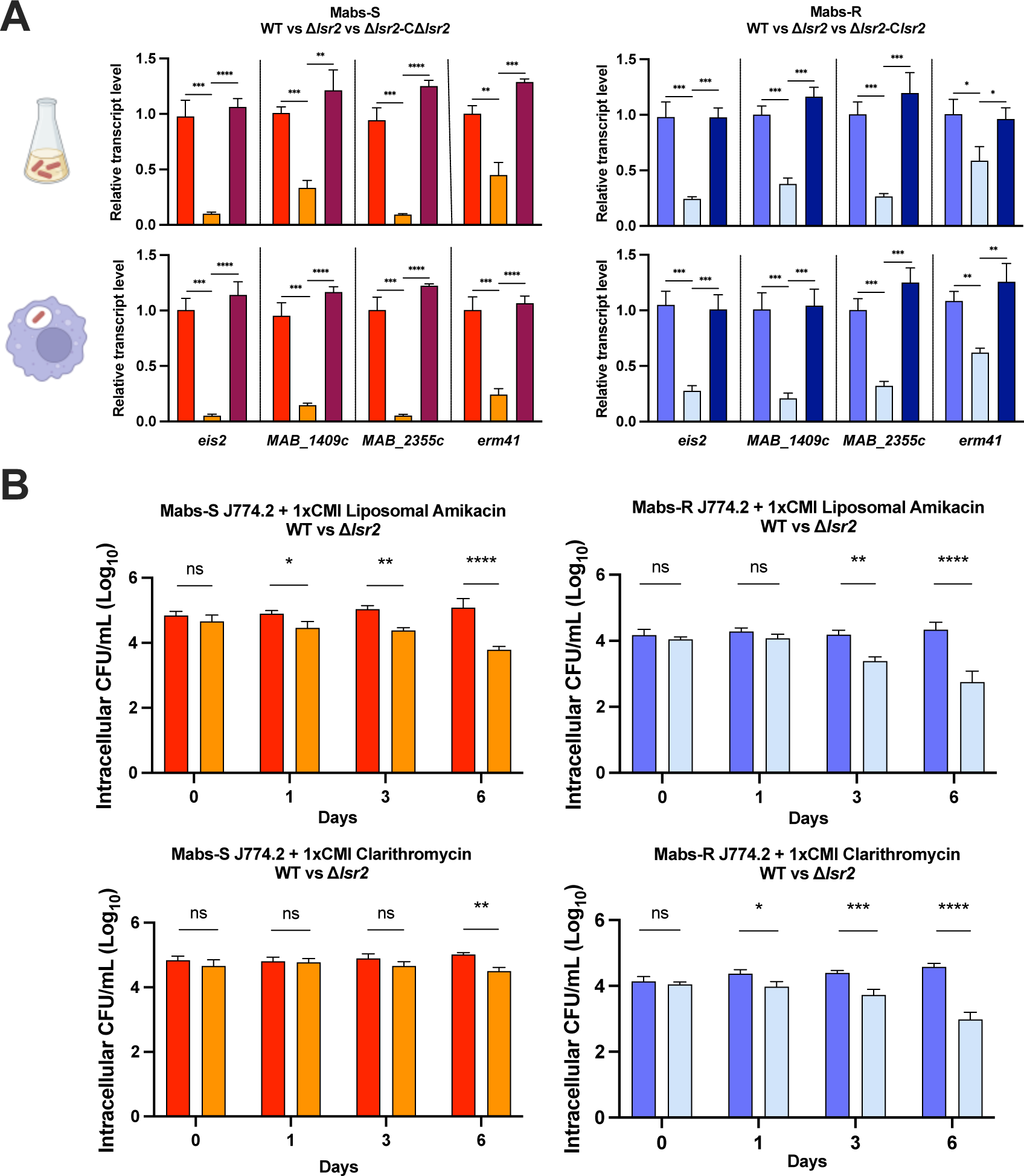
Enhanced sensitivity to aminoglycosides and macrolides due to diminished transcriptional expression of antimicrobial resistance genes in Mabs-S-Δ*lsr2* and Mabs-R-Δ*lsr2* strains. **A.** Quantification of relative expression of antimicrobial resistance genes; *eis2, MAB_1409c*, *MAB_2355c and erm41* by RT-qPCR under planktonic growth conditions (upper side) and intracellular growth conditions (lower side) in Mabs-S-WT (red), Mabs-S-Δ*lsr2* (orange), Mabs-S-Δ*lsr2*-C*lsr2* (brown), Mabs-R-WT (blue), Mabs-R-Δ*lsr2* (light blue) and Mabs-R-Δ*lsr2*-C*lsr2* (dark blue). *SigA* transcript levels were used for normalization. For each assay, n= 3 and error bars are SEM. *∗P < 0.05, ∗∗P < 0.01, ∗∗∗P < 0.001,* and *∗∗∗∗P < 0.0001* (Student’s *t* test). **B.** Intracellular growth of *M. abscessus* strains: Mabs-S-WT (red), Mabs-S-Δ*lsr2* (orange), Mabs-R-WT (blue), and Mabs-R-Δ*lsr2* (light blue), following antibiotic treatments. Murine J774.2 macrophages were infected with mycobacteria at an MOI of 10 and subsequently treated with 1X CMI liposomal amikacin or 1X CMI clarithromycin. Intracellular growth post-antibiotic treatment was assessed by counting CFUs at various time points after treatment (days 0, 1, 3, and 6). The data are representative of three independent experiments and are presented as means ± SEM. Differences between means were analyzed by two-way ANOVA and the Tukey post-test, allowing multiple comparisons. ns, non-significant, *∗P < 0.05, ∗∗P < 0.01, ∗∗∗P < 0.001,* and *∗∗∗∗P < 0.0001*.

**Table 1:** Increased sensitivity of *lsr2* mutant strains to aminoglycoside and macrolide antibiotics. Antibiotic susceptibility profiles (MICs) of *M. abscessus* wild-type (WT) and mutant (Δ*lsr2*) strains (μg/mL) after 3 to 5 days of incubation.

| Antimicrobial agent | | MIC (μg/mL) for | | | |
| --- | --- | --- | --- | --- | --- |
| | | Mabs-S-WT | Mabs-S-Δ*lsr2* | Mabs-R-WT | Mabs-R-Δ*lsr2* |
| Aminoglycosides | Amikacin | 9 | 4.5 | 15.5 | 8 |
|  | Tobramycin | 9.5 | 4.5 | 11 | 5.5 |
| Macrolides | Clarithromycin | 0.5 | 0.5 | 0.5 | 0.125 |

### DNA-binding profiles of Lsr2 on Mabs-S and Mabs-R genomes

We used ChIP-seq to reveal how Lsr2 binding mode on *M. abscessus* genome can explain its effect on transcription regulation. Lsr2 binds 6.9% (337/4920 genes) of *M. abscessus* genes, which is significantly less than what was observed in *M. tuberculosis* and *M. smegmatis* (respectively 21% and 13%) (20). Peak calling analysis showed that Lsr2 binding profile is similar between Mabs-S and Mabs-R with 348 enrichment peaks detected along the chromosome (**Figure 5A**). Lsr2 binds to long tracts of DNA, forming extensive binding domains that can span over 1 Kb and extend up to 6 Kb in certain regions (averages of 589 and 560 bp in Mabs-S and Mabs-R respectively) (**Figures 5B** and **5C**). This ability to form long binding domains shares similarities with other NAPs such as H-NS in Gram negative bacteria and has also been described for Lsr2 *in vivo* in *M. tuberculosis* (24, 54, 55). Furthermore, we found that Lsr2 binds preferentially to AT-rich sequences (43% on average) (**Figure 5B** and **5D, Supplementary figure S2)** compared to the average AT content observed in *M. abscessus* genome (35.9%) (7). Lsr2 had the same binding preference in other mycobacteria such as *M. tuberculosis* (20) and *M. smegmatis* (28). We next assess the distribution of Lsr2 over the different predicted operons constituting *M. abscessus’* genome. This analysis revealed that Lsr2 binds to 18% of the operons, with the ability to bind either operon promoters (12%), CDS (34%), or both operon promoters and CDS (54%) (**Figure 5E**). These findings emphasize the preferential enrichment of Lsr2 on coding sequences, suggesting its direct role in gene regulation through extensive binding domains.

**Figure 5.**
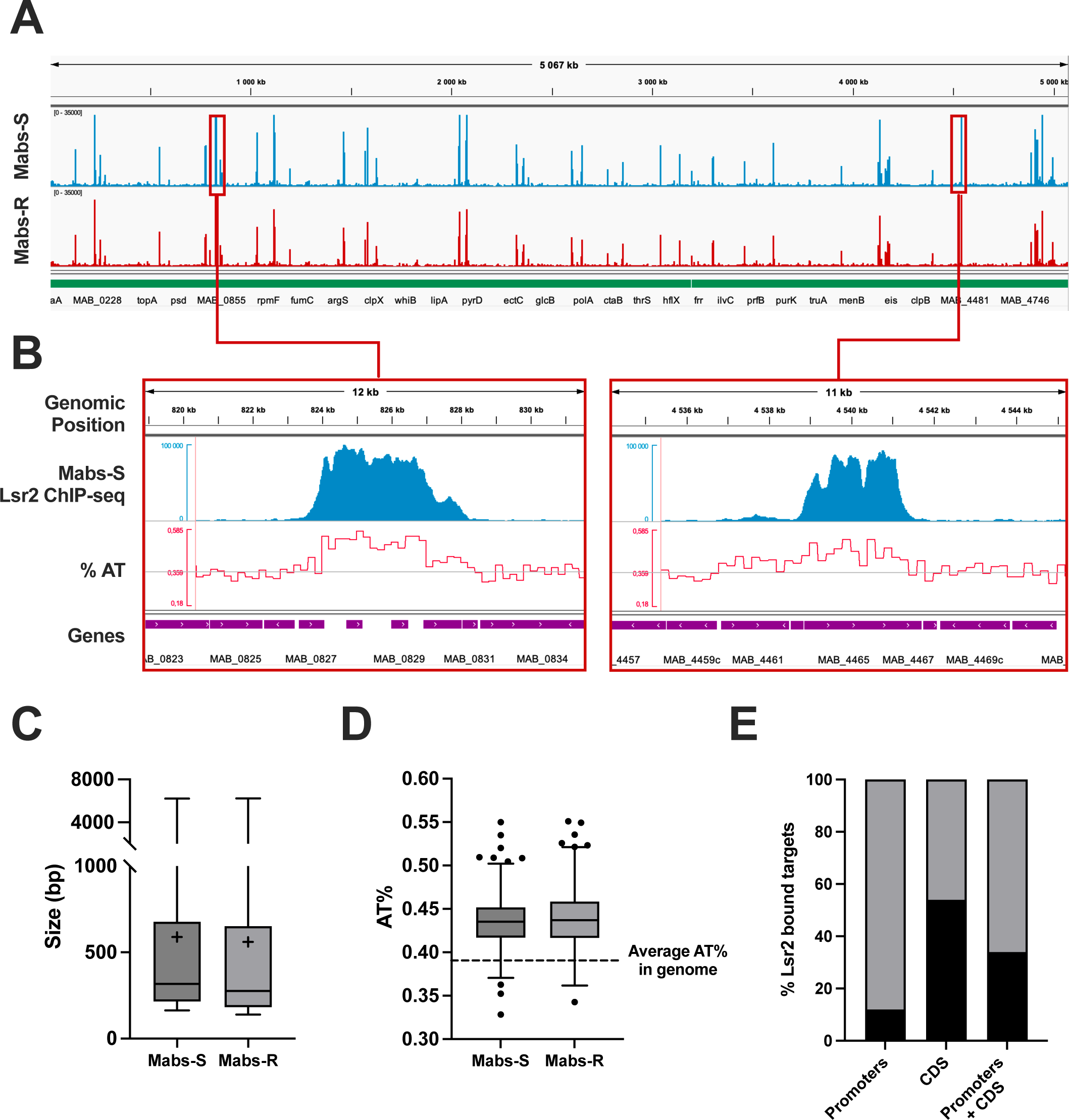
Modalities of Lsr2 binding on the *M. abscessus* genome. **A.** Global distribution of Lsr2 on the genomes of Mabs-S and Mabs-R morphotypes analyzed by ChIP-seq. **B**. Distribution of Lsr2 is strongly correlated with enhanced AT content as exemplified for genomic regions encompassing genes from *MAB_0827* to *MAB_0842* and from *MAB_4463* to *MAB_4467*. The bottom part corresponds to gene positions. C. Boxplot showing the distribution of the lengths of binding regions of Lsr2 in the Mabs-S and Mabs-R morphotypes (with mean indicated as a « + »). **D.** Boxplots summarizing the preference of Lsr2 to bind to AT-rich regions of the *M. abscessus* Mabs-S and Mabs-R genome. **E**. Histogram representing the percentage of Lsr2 within genomics features of operons (promoters, CDS or promoters with CDS).

### Effects of Lsr2 binding on transcription regulation in *M. abscessus*

Integration of RNA-seq data with Lsr2 binding profiles show that direct Lsr2 targets genes constitute only 29.8% and 41.3% of all Lsr2-regulated genes in Mabs-S and Mabs-R respectively, confirming the indirect mode of Lsr2 regulation. Secondly, although Lsr2 binds to a large number of genes, many of them remain unregulated, indicating that its binding to DNA alone is insufficient to trigger transcription regulation. Indeed, about 82% of the genes bound by Lsr2 are not differentially expressed in both Mabs-S and Mabs-R mutant (**Figure 6A**). If this lack of regulation can be explained by an absence of expression for a part of these genes in our experimental conditions, this result also suggests that unknown factors required for the transcriptional effect of Lsr2 are absent, or that Lsr2 has only a structural effect on these genomic regions. Furthermore, among all the direct targets of Lsr2 (**Figure 6B**), we found a stronger direct effect for Lsr2 in the regulation of genes involved in transport (69.7%) while all the antimicrobial resistance genes studied are indirectly regulated. To confirm the repressive function of Lsr2, we compared the expression levels between all Lsr2-regulated genes with its direct target genes in Mabs-S and Mabs-R. More than 78% of the direct target genes show significantly higher positive Log_2_FC values compared to all Lsr2-regulated genes in both morphotypes, confirming the direct repressive role of Lsr2 in gene regulation (**Figure 6C**). In order to assess the potential impact of the length of Lsr2 binding regions on its repressive effect, we investigated the distribution of expression levels among the direct target genes, categorized based on the size of the binding domains formed by Lsr2 (either less than 1 Kb or greater than 1 Kb). Significantly, more positive Log_2_FC folds are correlated with binding domains larger than 1 Kb (*p < 0.0001* in Mabs-S and *p < 0.01* in Mabs-R), suggesting a positive correlation between the size of Lsr2-enriched domains and the repressive effect of Lsr2 (**Figure 6D**).

**Figure 6.**
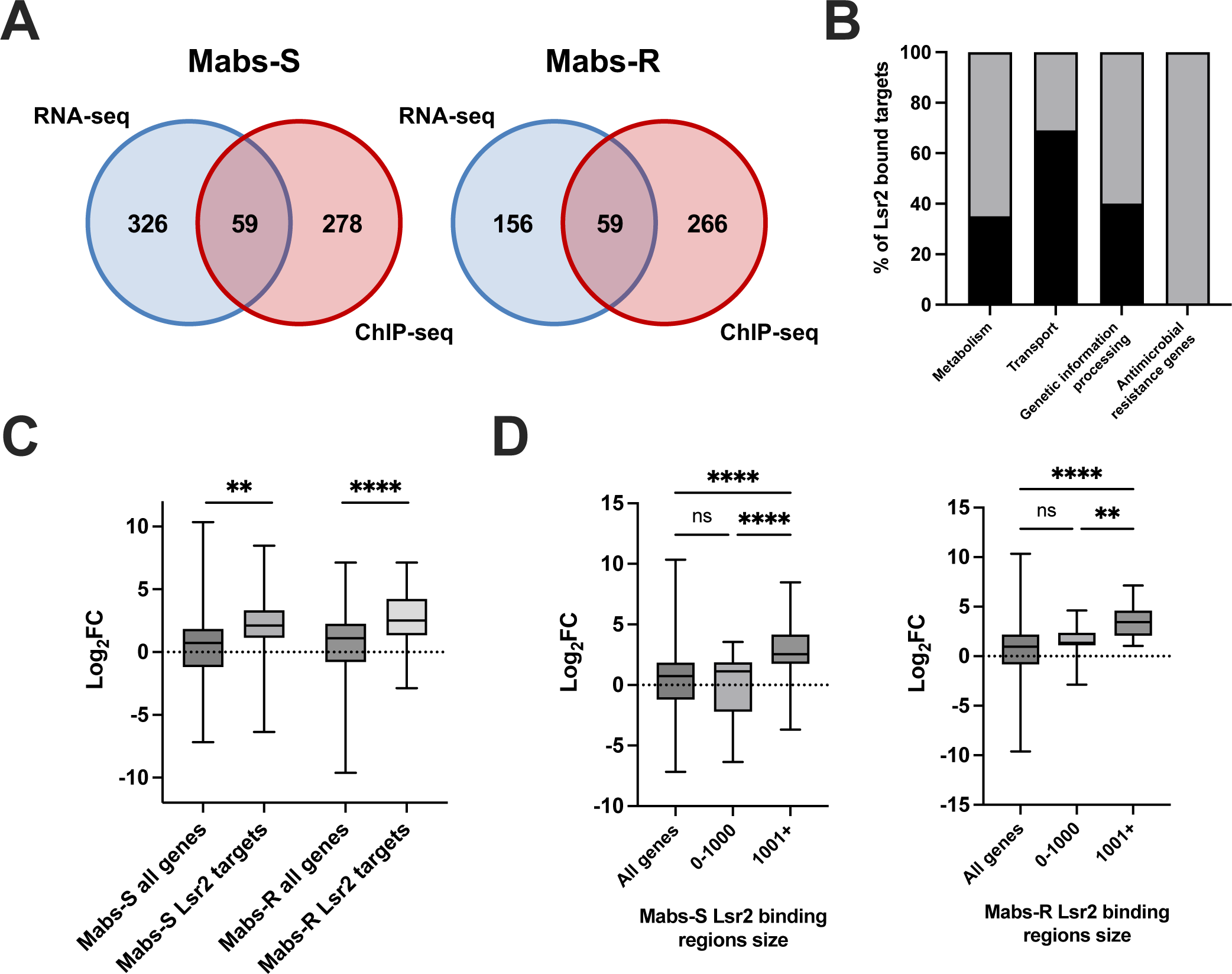
Integrative analysis RNA-seq/ChIP-seq. **A.** Venn diagram showing the number of direct target genes of Lsr2, the number of genes regulated by Lsr2 without binding and the number of genes binding by Lsr2 without regulatory effect in Mabs-S and Mabs-R. **B.** Histogram representing the percentage of direct target genes of Lsr2 in the categories of metabolism, transport, genetic information processing and antimicrobial resistance genes. **C**. Box plots comparing Log_2_FC of all genes regulated by Lsr2 to Log_2_FC of direct target genes in Mabs-S and Mabs-R. **D**. Box plots comparing Log_2_FC of all genes regulated by Lsr2 to Log_2_FC of direct target genes belonging to two domain size categories formed by Lsr2 less than 1 Kb and greater than 1 Kb. Differences between means were analyzed by ordinary one-way ANOVA. ns, non-significant, *∗∗P < 0.01,* and *∗∗∗∗P < 0.0001*.

## DISCUSSION

If NAPs are widely recognized as essential factors in the hierarchical structural organization of the bacterial chromosome, compelling evidence also highlights their crucial role in pathogen adaptation to changing environments, including their ability to thrive within host cells, such as macrophages, during infection. This includes reactive oxygen species (ROS) production, pH variations, hypoxia or nutrient scarcity (56). The present study further establishes the pivotal role of the NAP Lsr2 in the infectious strategy of *M. abscessus* as well as in its antibiotic resistance. Consistent with previous findings (29), our transcriptomic analysis confirms the impact of Lsr2 in controlling expression of key virulence factors on the intracellular and *in vivo* survival of *M. abscessus*. Among *lsr2* regulated genes, *eis2* is an important determinant of intracellular growth whose expression is highly induced in macrophages during infection. It controls the production of ROS, the sensitivity of bacteria to hydrogen peroxide (H_2_O_2_) and contributes to phagosomal membrane damage, thereby facilitating contact between the pathogen and cytosol (30). Similarly, *paa* genes that are upregulated by *lsr2* contribute to stress tolerance induced by antibiotics in *Acinetobacter baumannii* (57). These genes are also present in *Burkholderia cepacia* where the *paak*’s homolog is essential for the survival of the pathogen during infection in a rat model of chronic lung (58). The *sigD* and *SigE* genes, encoding two sigma factors regulated by Lsr2 in Mabs-S and Mabs-R respectively, are involved in macrophage viability through resistance to oxidative stress. The loss of *sigD* reduces virulence, histopathology and lethality of *M. tuberculosis* infection in mice (36–38). The KEGG analysis highlights a role of Lsr2 in the regulation of multiple metabolic pathways, with a particular emphasis for genes related to fatty acid degradation. This is intriguing, considering that during macrophage infections, a metabolic shift in carbon source utilization from sugars to lipids as a carbon source was previously observed (30), reflecting certain signatures of adaptation in intracellular life. Hence, the observed increase in mortality among *lsr2* mutant strains during infection (29) may be attributed to the challenges they face in adapting their metabolism in the absence of Lsr2. Likewise, Fe-S proteins, also regulated by Lsr2, are intracellular stress sensors and play a role in the genomic stability whose biosynthesis was induced in macrophages and amoeba by *M. abscessus* to adapt to an intracellular lifestyle (30). It has also been reported that impaired Fe-S protein biogenesis was deleterious to *Pseudomonas aeruginosa* growth (59) and decreases the ability of *Shigella flexneri* to invade epithelial cells lining the human colon (60).

Treatment of *M. abscessus* infection is particularly challenging, as this bacterium is resistant to most classes of antibiotics (9, 61). Here, our transcriptomic analysis revealed that four genes (*eis2, MAB_1409c, MAB_2355c* and *erm41*) contributing to aminoglycoside and macrolide resistance, are regulatory targets activated by Lsr2 in both morphotypes. *erm41,* encoding a methyltransferase, is widely studied for its role in inducible macrolide resistance in *M. abscessus,* observed in 40-60% of clinical strains (49, 50, 62). *eis2*, which code for an N-acetyltransferase, plays an important role in aminoglycoside resistance in *M. abscessus* by acetylating the first amino group of molecules like amikacin, in addition to its role in intracellular growth (53). The role of Lsr2 as a modulator of antibiotic resistance has been previously demonstrated in mycobacteria (63). In *M. smegmatis*, deletion of *lsr2* showed increased sensitivity of cells to nalidixic acid and rifampicin compared to the wild-type strain, in addition to better penetration for vancomycin through the cell wall (28). Similarly, in *M. abscessus*, we reported a greater sensitivity of *lsr2* mutant strains to liposomal amikacin and clarithromycin during macrophage infections and MICs determination. The increased sensitivity to antibiotics following *lsr2* deletion could also be explained by a deregulation of certain enzymes involved in the metabolism of lipids essential for cell wall formation, thereby modifying its permeability. Alternatively, the contribution of Lsr2 to antibiotic resistance in *M. tuberculosis* has been explained by the repression of an efflux pump for isoniazid following its overexpression (63). However, according to ChIP-seq results, these genes show no Lsr2 enrichment, suggesting their indirect regulation. Overall, the regulatory role of Lsr2 protein in response to antibiotics across various mycobacteria confirmed its potential as a target for the development of new antimicrobials.

Lsr2 was previously shown to bind to the promoter of the *mps* operon, while its deletion increases GPL level at the membrane of *M. smegmatis* (24, 47). However, we previously showed that the *lsr2* mutation did not restore the production of GPL in Mabs-R (29). These findings align with the identification of a CG insertion in the 5’ region of the *mps1* gene, which completely abolishes the production of transcripts for the *mps1-mps2-gap* operon in this morphotype (17). Remarkably, our RNA-seq data demonstrated that *lsr2* deletion can restore transcripts elongation in this region to levels comparable to those observed in Mabs-S. Nevertheless, the presence of this indel is likely to result in a non-functional protein, thus preventing the bacteria to transition back to a smooth morphotype. This result still suggests an intriguing insight regarding Lsr2 functions in regulating transcription dynamics and RNA polymerase processivity, possibly via oligomerization or through changes in local DNA topology.

One of the fundamental goals of our study was to highlight the binding modes and to identify the direct target genes of Lsr2 on Mabs-S and Mabs-R genomes. Our ChIP-seq analysis identified 348 Lsr2 binding sites distributed throughout the *M. abscessus* chromosome. Like its ortholog H-NS, we have determined that Lsr2 presents a high affinity for AT-rich sequences. Previous reports suggest that Lsr2 and H-NS act as xenogeneic silencers for AT-rich regions acquired by horizontal gene transfer (21, 64, 65). In the case of *M. abscessus*, our findings do not support the binding of Lsr2 to these specific genes. Analysis of the Lsr2 genomic distribution in *M. tuberculosis* showed an enrichment pattern across 840 genes, including genes coding for type VII secretion systems of the ESX family, for the synthesis of cell wall lipids and PE/PPE proteins. In contrast, neither our ChIP-seq nor RNA-seq data revealed any binding or regulation by Lsr2 on genes involved in the secretion systems in *M. abscessus*. Our data also show that Lsr2 binding spans genomic regions that could sometimes exceed 1 Kb in size. As already suggested in previous works, this observation highlights the potential role of Lsr2 in repressing gene expression through a mechanism involving its oligomerization along promoters and coding sequences of its target genes. It has also been suggested that the oligomerization of Lsr2 plays a significant role in enhancing DNA stability by safeguarding it against nuclease degradation or ROS (66).

Finally, the existence of a limited repertoire of shared Lsr2-regulated genes between Mabs-S and Mabs-R firstly suggests differences in the Lsr2 distribution on the two morphotypes. Surprisingly, this distribution was strictly identical between both morphotypes with the same number of enrichment peaks detected. This result was further confirmed by a quantitative statistical analysis using the DiffBind R package. It has been reported that the mode of action of Lsr2 and H-NS could depend on additional features, such as the bridging of oligomers on distant DNA fragments, which most likely contribute to loop formation (24, 25). The different regulation between the two morphotypes as well as the presence of a large number of genes bound by Lsr2 without any effect on their transcription strongly suggests the intervention of other factors such as the existence of partner proteins or post-transcriptional modifications to explain the Lsr2-dependant transcriptional repression mechanism.

## MATERIALS AND METHODS

### *M. abscessus* strains, growth conditions and DNA manipulations

Wild-type, *lsr2* mutant and *lsr2* complemented strains of *Mycobacterium abscessus* 19977-IP, designated Mabs-S-WT, Mabs-R-WT, Mabs-S-Δ*lsr2*, Mabs-R-Δ*lsr2,* Mabs-S-Δ*lsr2*-C*lsr2* and Mabs-R-Δ*lsr2*-C*lsr2,* were grown aerobically at 37°C in Middlebrook 7H9 broth or on Middlebrook 7H11 agar, supplemented with 2% glycerol, 1% glucose and 0.05% tween 80 when necessary. Construction of *lsr2* mutant and *lsr2* complemented strains was carried out during our previous study (29). Zeocin (25 µg/mL), kanamycin (250 µg/mL) and hygromycin (500 µg/mL) were added to cultures for strain selection. The Lsr2-FLAG expressing strains used for ChIP-seq analysis, designated Mabs-S-Lsr2-FLAG and Mabs-R-Lsr2-FLAG were obtained by cloning PCR-amplified homology regions surrounding the *lsr2* locus into a vector containing the *lsr2* gene sequence fused with three repeats of the FLAG epitope followed with a zeocin resistance cassette. Sequences of primers used for this construction are listed in **Supplementary Table S1**. Replacement of the endogenous *lsr2* gene by *lsr2*-FLAG-zeocin sequence in Mabs-S and Mabs-R was performed as described previously (28, 55, 67). The insertion of the FLAG protein did not alter the phenotype of the bacteria, nor did it affect the *in vitro* growth rates of the strains in liquid culture when compared to Mabs-S and Mabs-R (**Supplementary Figure S3**).

### RNA extraction, purification and library preparation for RNA sequencing

Wild-type and *lsr2* mutant strains were grown until exponential growth phase with optical density value 600 nm (OD_600_) of 0.6 corresponding to approximately 2.4 × 10^8^ cells/mL. A total of 4.8 × 10^9^ cells were harvested and total RNA from three biological replicates was extracted for each strain using the Monarch® Total RNA Miniprep (NEB) Kit. After DNase I treatment and elution, RNA concentrations were measured using the Qubit fluorometer (Promega). RNA integrity was evaluated using the Bioanalyzer system (Agilent) and all samples used for library preparation showed RNA Integrity Number (RIN) scores >8. rRNA depletion was performed using the NEBNext® rRNA Depletion Kit (Bacteria) (NEB) and the sequencing libraries were prepared using NEBNext Ultra II Directional RNA Library Prep Kit (NEB) following manufacturer’s instructions. RNA extraction from *lsr2* complemented strains was carried out in the same for RT-qPCR experiments.

### NGS sequencing and data analyses

All libraries were sequenced using a NextSeq500 (Illumina) at the Genomics platform of the UFR Sciences de la Santé (UVSQ, Montigny-le-Bretonneux, France). The reads were trimmed with fastp v0.23.2 (68) and mapped to the *M. abscessus* ATCC 19977 reference genome (NCBI accession number NC_010397) using STAR 2.7.6a (69). Reads were counted with featureCounts v2.0 (70). Differential gene expression analysis was performed on R using the SARtools package using correction for batch effect (71). Significantly activated/repressed genes were selected with a threshold of 0.05 on the adjusted P-value and a Log_2_ fold change (Log_2_FC) of at least 1.

### Gene Ontology term analysis

The Gene Ontology (GO) annotation of *M. abscessus* genome, specifically BP (Biological Process) and MF (Molecular Function) categories, was used to describe the roles and functions of the differentially expressed genes. The enrichment analysis was performed using the R package TopGO v.2.46.0 by combining the three different methods (*classic*, *elim* and *weight*) with two statistical tests to evaluate the score significance (Fisher’s exact test and Kolmogorov-Smirnov test).

### ChIP-seq library preparation

Mabs-S-Lsr2-FLAG and Mabs-R-Lsr2-FLAG strains were grown until an OD of 0.6 and ChIP experiments were carried out in three biological replicates for each strain. Briefly, 50 mL of culture was fixed in 1% formaldehyde (Euromedex), quenched with glycine, washed with phosphate-buffered saline (PBS) and lysed using VK05 beads and Precellys grinder (3 cycles: 8,700 rpm −3 × 20 s ON/60 s OFF, Bertin Technologies). For immunoprecipitation, 25 µg of chromatin was incubated with 25 µl of monoclonal anti-FLAG® M2 magnetic beads (Sigma-Aldrich) on a rotary shaker for 16 h at 4°C (29). The immunoprecipitated DNA and 1% of the total input were reverse-crosslinked and eluted using the iPure v2 kit (Diagenode). For library preparation, NEBNext® Ultra™ II DNA Library Prep Kit for Illumina (NEB) was used. In parallel, ChIP-Seq libraries of Mabs-S and Mabs-R wild-type strains were prepared as a negative control. All the libraries were sequenced with paired ends (150 cycles) using an iSeq100 sequencer (Illumina).

### ChIP-seq data analysis

Reads were trimmed with fastp v0.23.2 (68), followed by mapping to the *M. abscessus* ATCC 19977 genome (NCBI accession number NC_010397) using Bowtie v2.4.5 (72) with a distribution of read coverages showing a mean of 35 to 58 as a proxy for sequencing depth. Fragments size of all libraires showed a distribution with median values ranging from 107 to 165 bp. PCR duplicates were marked by Picard 3.1.0 and further removed with samtools v1.15.1 (73). Peaks were visualized as raw sequencing coverage using the IGV genome browser (74) after conversion of the bam files in bigwig using Deeptools (75) without input normalization. Peak calling was performed using MACS v2.1.3.3 (76) with an FDR = 0,0029 and normalized against input libraries. Occurrence of Lsr2 peaks on operons or their promoters was detected using the intersect function of Bedtools v2.29.2 (77). The *M. abscessus* operons were predicted using Operon-mapper (78). Differential analysis of Lsr2 peaks between Mabs-S and Mabs-R was performed on R using the DiffBind package v3.10.0 (79).

### Drug susceptibility testing

Minimum inhibitory concentrations (MICs) were determined for amikacin (1-256 µg/mL), tobramycin (0.25-64 µg/mL), clarithromycin (0.06-32 µg/mL) according to Clinical & Laboratory Standards Institute (CLSI) guidelines (80). Stock solutions of amikacin (Mylan) and tobramycin (Mylan) were dissolved in water and stock solutions of clarithromycin (Mylan) were dissolved in dimethyl sulfoxide (DMSO). Drug susceptibility testing was determined using the microdilution method, cation-adjusted Mueller-Hinton broth in three biological replicates, as described previously (81, 82).

### Macrophage culture, infections and intracellular survival after antibiotic treatments

Murine J774.2 macrophages were maintained at 37°C under 5% CO2 in Dulbecco’s modified Eagle medium (DMEM) (Gibco) supplemented with 10% fetal bovine serum (FBS) (Gibco), penicillin (100 IU/mL) and streptomycin (100 µg/mL). Macrophages were seeded in a 24-well plate at a concentration of 5×10^4^ cells/mL of medium. Macrophage infections with wild-type and *lsr2* mutant strains at 10 MOI and treatments with 1xCMI liposomal amikacin (Insmed) and clarithromycin (Mylan) were carried out in three biological replicates, as described previously (16, 83, 84). The number of intracellular CFU/mL was determined on days 1, 3, and 6 post-infection and post-treatment. For plate day 0, the number of intracellular CFU/mL was determined 3 hours after infection, without any antibiotic treatments. In parallel, the same infections were performed without antibiotic treatments as control for bacterial growth for 3 plates (days 1, 3 and 6).

### RNA extraction of mycobacteria from macrophage cocultures

Infections and RNA extractions were carried out in three biological replicates as described previously (30, 83). In brief, approximately 10^7^ cells of murine J774.2 macrophages were infected with wild-type, *lsr2* mutant and *lsr2* complemented strains at 50 MOI. Macrophages were lysed 16 hours post-infection with a guanidine thiocyanate solution (4 M) plus β-mercaptoethanol and total RNA was isolated from the bacterial pellets using TRIzol reagent (Ambion) in the presence of zirconia/silica beads. After bacterial cell disruption using a Precellys grinder (3 cycles: 8,700 rpm −3 × 20 s ON/60 s OFF, Bertin Technologies), RNA samples were precipitated using chloroform isoamyl alcohol 24:1 and with 0.8 volumes of cold isopropanol, washed with ethanol (70%), re-suspended in RNase-free water and treated with the Turbo DNA-free kit (Invitrogen) to remove DNA contaminants.

### Reverse transcriptase quantitative PCR (RT-qPCR)

cDNA was prepared using SuperScript® III First-Strand Synthesis System Kit (Invitrogen). RT-qPCR was performed with a CFX96 thermal cycler (Bio-Rad), using the DyNAmo ColorFlash SYBR Green qPCR Kit (ThermoFisher), as described previously (14, 17). Target gene expression was quantified relative to the reference gene, *sigA*. The sequences of primers used for quantitative real-time PCR (RT-qPCR) are listed in **Supplementary Table S1**. Each RT-qPCR was performed on three biological replicates.

## DATA AVAILABILITY

All genomic data produced in the present project (RNA-seq and ChIP-seq) are available in the NCBI GEO database under accession number GSE239871.

## Author contributions

E.G. designed and performed experiments, analyzed data, and wrote the paper; M.P.R-O. analyzed data; ND wrote the paper; J-L.H. designed experiments and wrote the paper; F.C. designed experiments, analyzed data, and wrote the paper.

## REFERENCES

1. Thomson RM, Carter R, Tolson C, Coulter C, Huygens F, Hargreaves M. 2013. Factors associated with the isolation of Nontuberculous mycobacteria (NTM) from a large municipal water system in Brisbane, Australia. BMC Microbiol 13:89.

2. Mougari F, Guglielmetti L, Raskine L, Sermet-Gaudelus I, Veziris N, Cambau E. 2016. Infections caused by Mycobacterium abscessus: epidemiology, diagnostic tools and treatment. Expert Rev Anti Infect Ther 14:1139–1154.

3. Degiacomi G, Sammartino JC, Chiarelli LR, Riabova O, Makarov V, Pasca MR. 2019. Mycobacterium abscessus, an Emerging and Worrisome Pathogen among Cystic Fibrosis Patients. Int J Mol Sci 20:5868.

4. Johansen MD, Herrmann J-L, Kremer L. 2020. Non-tuberculous mycobacteria and the rise of Mycobacterium abscessus. Nat Rev Microbiol 18:392–407.

5. Qvist T, Taylor-Robinson D, Waldmann E, Olesen HV, Hansen CR, Mathiesen IH, Høiby N, Katzenstein TL, Smyth RL, Diggle PJ, Pressler T. 2016. Comparing the harmful effects of nontuberculous mycobacteria and Gram negative bacteria on lung function in patients with cystic fibrosis. J Cyst Fibros Off J Eur Cyst Fibros Soc 15:380–385.

6. Esther CR, Esserman DA, Gilligan P, Kerr A, Noone PG. 2010. Chronic Mycobacterium abscessus infection and lung function decline in cystic fibrosis. J Cyst Fibros Off J Eur Cyst Fibros Soc 9:117–123.

7. Ripoll F, Pasek S, Schenowitz C, Dossat C, Barbe V, Rottman M, Macheras E, Heym B, Herrmann J-L, Daffé M, Brosch R, Risler J-L, Gaillard J-L. 2009. Non mycobacterial virulence genes in the genome of the emerging pathogen Mycobacterium abscessus. PloS One 4:e5660.

8. Rominski A, Selchow P, Becker K, Brülle JK, Dal Molin M, Sander P. 2017. Elucidation of Mycobacterium abscessus aminoglycoside and capreomycin resistance by targeted deletion of three putative resistance genes. J Antimicrob Chemother 72:2191–2200.

9. Illouz M, Alcaraz M, Roquet-Banères F, Kremer L. 2021. [Mycobacterium abscessus, a model of resistance to multiple antibiotic classes]. Med Sci MS 37:993–1001.

10. Byrd TF, Lyons CR. 1999. Preliminary characterization of a Mycobacterium abscessus mutant in human and murine models of infection. Infect Immun 67:4700–4707.

11. Howard ST, Rhoades E, Recht J, Pang X, Alsup A, Kolter R, Lyons CR, Byrd TF. 2006. Spontaneous reversion of Mycobacterium abscessus from a smooth to a rough morphotype is associated with reduced expression of glycopeptidolipid and reacquisition of an invasive phenotype. Microbiol Read Engl 152:1581–1590.

12. Medjahed H, Gaillard J-L, Reyrat J-M. 2010. Mycobacterium abscessus: a new player in the mycobacterial field. Trends Microbiol 18:117–123.

13. Davidson LB, Nessar R, Kempaiah P, Perkins DJ, Byrd TF. 2011. Mycobacterium abscessus glycopeptidolipid prevents respiratory epithelial TLR2 signaling as measured by HβD2 gene expression and IL-8 release. PloS One 6:e29148.

14. Roux A-L, Ray A, Pawlik A, Medjahed H, Etienne G, Rottman M, Catherinot E, Coppée J-Y, Chaoui K, Monsarrat B, Toubert A, Daffé M, Puzo G, Gaillard J-L, Brosch R, Dulphy N, Nigou J, Herrmann J-L. 2011. Overexpression of proinflammatory TLR-2-signalling lipoproteins in hypervirulent mycobacterial variants. Cell Microbiol 13:692–704.

15. Bernut A, Herrmann J-L, Kissa K, Dubremetz J-F, Gaillard J-L, Lutfalla G, Kremer L. 2014. Mycobacterium abscessus cording prevents phagocytosis and promotes abscess formation. Proc Natl Acad Sci U S A 111:E943–952.

16. Roux A-L, Viljoen A, Bah A, Simeone R, Bernut A, Laencina L, Deramaudt T, Rottman M, Gaillard J-L, Majlessi L, Brosch R, Girard-Misguich F, Vergne I, de Chastellier C, Kremer L, Herrmann J-L. 2016. The distinct fate of smooth and rough Mycobacterium abscessus variants inside macrophages. Open Biol 6:160185.

17. Pawlik A, Garnier G, Orgeur M, Tong P, Lohan A, Le Chevalier F, Sapriel G, Roux A-L, Conlon K, Honoré N, Dillies M-A, Ma L, Bouchier C, Coppée J-Y, Gaillard J-L, Gordon SV, Loftus B, Brosch R, Herrmann JL. 2013. Identification and characterization of the genetic changes responsible for the characteristic smooth-to-rough morphotype alterations of clinically persistent Mycobacterium abscessus. Mol Microbiol 90:612–629.

18. Laal S, Sharma YD, Prasad HK, Murtaza A, Singh S, Tangri S, Misra RS, Nath I. 1991. Recombinant fusion protein identified by lepromatous sera mimics native Mycobacterium leprae in T-cell responses across the leprosy spectrum. Proc Natl Acad Sci U S A 88:1054–1058.

19. Chen JM, German GJ, Alexander DC, Ren H, Tan T, Liu J. 2006. Roles of Lsr2 in colony morphology and biofilm formation of Mycobacterium smegmatis. J Bacteriol 188:633–641.

20. Gordon BRG, Li Y, Wang L, Sintsova A, van Bakel H, Tian S, Navarre WW, Xia B, Liu J. 2010. Lsr2 is a nucleoid-associated protein that targets AT-rich sequences and virulence genes in Mycobacterium tuberculosis. Proc Natl Acad Sci U S A 107:5154–5159.

21. Gehrke EJ, Zhang X, Pimentel-Elardo SM, Johnson AR, Rees CA, Jones SE, Hindra null, Gehrke SS, Turvey S, Boursalie S, Hill JE, Carlson EE, Nodwell JR, Elliot MA. 2019. Silencing cryptic specialized metabolism in Streptomyces by the nucleoid-associated protein Lsr2. eLife 8:e47691.

22. Dame RT, Rashid F-ZM, Grainger DC. 2020. Chromosome organization in bacteria: mechanistic insights into genome structure and function. Nat Rev Genet 21:227–242.

23. Hołówka J, Zakrzewska-Czerwińska J. 2020. Nucleoid Associated Proteins: The Small Organizers That Help to Cope With Stress. Front Microbiol 11:590.

24. Chen JM, Ren H, Shaw JE, Wang YJ, Li M, Leung AS, Tran V, Berbenetz NM, Kocíncová D, Yip CM, Reyrat J-M, Liu J. 2008. Lsr2 of Mycobacterium tuberculosis is a DNA-bridging protein. Nucleic Acids Res 36:2123–2135.

25. Liu J, Gordon BRG. 2012. Targeting the global regulator Lsr2 as a novel approach for anti-tuberculosis drug development. Expert Rev Anti Infect Ther 10:1049–1053.

26. Colangeli R, Haq A, Arcus VL, Summers E, Magliozzo RS, McBride A, Mitra AK, Radjainia M, Khajo A, Jacobs WR, Salgame P, Alland D. 2009. The multifunctional histone-like protein Lsr2 protects mycobacteria against reactive oxygen intermediates. Proc Natl Acad Sci U S A 106:4414–4418.

27. Bartek IL, Woolhiser LK, Baughn AD, Basaraba RJ, Jacobs WR, Lenaerts AJ, Voskuil MI. 2014. Mycobacterium tuberculosis Lsr2 is a global transcriptional regulator required for adaptation to changing oxygen levels and virulence. mBio 5:e01106–01114.

28. Kołodziej M, Łebkowski T, Płociński P, Hołówka J, Paściak M, Wojtaś B, Bury K, Konieczny I, Dziadek J, Zakrzewska-Czerwińska J. 2021. Lsr2 and Its Novel Paralogue Mediate the Adjustment of Mycobacterium smegmatis to Unfavorable Environmental Conditions. mSphere 6:e00290–21.

29. Le Moigne V, Bernut A, Cortès M, Viljoen A, Dupont C, Pawlik A, Gaillard J-L, Misguich F, Crémazy F, Kremer L, Herrmann J-L. 2019. Lsr2 Is an Important Determinant of Intracellular Growth and Virulence in Mycobacterium abscessus. Front Microbiol 10:905.

30. Dubois V, Pawlik A, Bories A, Le Moigne V, Sismeiro O, Legendre R, Varet H, Rodríguez-Ordóñez MDP, Gaillard J-L, Coppée J-Y, Brosch R, Herrmann J-L, Girard-Misguich F. 2019. Mycobacterium abscessus virulence traits unraveled by transcriptomic profiling in amoeba and macrophages. PLoS Pathog 15:e1008069.

31. Alsarraf HMAB, Ung KL, Johansen MD, Dimon J, Olieric V, Kremer L, Blaise M. 2022. Biochemical, structural, and functional studies reveal that MAB_4324c from Mycobacterium abscessus is an active tandem repeat N-acetyltransferase. FEBS Lett 596:1516–1532.

32. El-Shazly S, Ahmad S, Mustafa AS, Al-Attiyah R, Krajci D. 2007. Internalization by HeLa cells of latex beads coated with mammalian cell entry (Mce) proteins encoded by the mce3 operon of Mycobacterium tuberculosis. J Med Microbiol 56:1145–1151.

33. Stavrum R, Valvatne H, Stavrum A-K, Riley LW, Ulvestad E, Jonassen I, Doherty TM, Grewal HMS. 2012. Mycobacterium tuberculosis Mce1 protein complex initiates rapid induction of transcription of genes involved in substrate trafficking. Genes Immun 13:496– 502.

34. Velmurugan K, Chen B, Miller JL, Azogue S, Gurses S, Hsu T, Glickman M, Jacobs WR, Porcelli SA, Briken V. 2007. Mycobacterium tuberculosis nuoG is a virulence gene that inhibits apoptosis of infected host cells. PLoS Pathog 3:e110.

35. Pradhan G, Shrivastva R, Mukhopadhyay S. 2018. Mycobacterial PknG Targets the Rab7l1 Signaling Pathway To Inhibit Phagosome-Lysosome Fusion. J Immunol Baltim Md 1950 201:1421–1433.

36. Calamita H, Ko C, Tyagi S, Yoshimatsu T, Morrison NE, Bishai WR. 2005. The Mycobacterium tuberculosis SigD sigma factor controls the expression of ribosome-associated gene products in stationary phase and is required for full virulence. Cell Microbiol 7:233–244.

37. Manganelli R. 2014. Sigma Factors: Key Molecules in Mycobacterium tuberculosis Physiology and Virulence. Microbiol Spectr 2:2.1.02.

38. Sachdeva P, Misra R, Tyagi AK, Singh Y. 2010. The sigma factors of Mycobacterium tuberculosis: regulation of the regulators. FEBS J 277:605–626.

39. Chen T, He L, Deng W, Xie J. 2013. The Mycobacterium DosR regulon structure and diversity revealed by comparative genomic analysis. J Cell Biochem 114:1–6.

40. Gautam US, Sikri K, Vashist A, Singh V, Tyagi JS. 2014. Essentiality of DevR/DosR interaction with SigA for the dormancy survival program in Mycobacterium tuberculosis. J Bacteriol 196:790–799.

41. Singh A, Crossman DK, Mai D, Guidry L, Voskuil MI, Renfrow MB, Steyn AJC. 2009. Mycobacterium tuberculosis WhiB3 maintains redox homeostasis by regulating virulence lipid anabolism to modulate macrophage response. PLoS Pathog 5:e1000545.

42. Mehta M, Rajmani RS, Singh A. 2016. Mycobacterium tuberculosis WhiB3 Responds to Vacuolar pH-induced Changes in Mycothiol Redox Potential to Modulate Phagosomal Maturation and Virulence. J Biol Chem 291:2888–2903.

43. You D, Xu Y, Yin B-C, Ye B-C. 2019. Nitrogen Regulator GlnR Controls Redox Sensing and Lipids Anabolism by Directly Activating the whiB3 in Mycobacterium smegmatis. Front Microbiol 10:74.

44. Viljoen A, Dubois V, Girard-Misguich F, Blaise M, Herrmann J-L, Kremer L. 2017. The diverse family of MmpL transporters in mycobacteria: from regulation to antimicrobial developments. Mol Microbiol 104:889–904.

45. Richard M, Gutiérrez AV, Viljoen A, Rodriguez-Rincon D, Roquet-Baneres F, Blaise M, Everall I, Parkhill J, Floto RA, Kremer L. 2019. Mutations in the MAB_2299c TetR Regulator Confer Cross-Resistance to Clofazimine and Bedaquiline in Mycobacterium abscessus. Antimicrob Agents Chemother 63:e01316–18.

46. Dubois V, Viljoen A, Laencina L, Le Moigne V, Bernut A, Dubar F, Blaise M, Gaillard J-L, Guérardel Y, Kremer L, Herrmann J-L, Girard-Misguich F. 2018. MmpL8MAB controls Mycobacterium abscessus virulence and production of a previously unknown glycolipid family. Proc Natl Acad Sci U S A 115:E10147–E10156.

47. Kocíncová D, Singh AK, Beretti J-L, Ren H, Euphrasie D, Liu J, Daffé M, Etienne G, Reyrat J-M. 2008. Spontaneous transposition of IS1096 or ISMsm3 leads to glycopeptidolipid overproduction and affects surface properties in Mycobacterium smegmatis. Tuberc Edinb Scotl 88:390–398.

48. Guo Q, Zhang Y, Fan J, Zhang H, Zhang Z, Li B, Chu H. 2021. MAB_2355c Confers Macrolide Resistance in Mycobacterium abscessus by Ribosome Protection. Antimicrob Agents Chemother 65:e0033021.

49. Nash KA, Brown-Elliott BA, Wallace RJ. 2009. A novel gene, erm(41), confers inducible macrolide resistance to clinical isolates of Mycobacterium abscessus but is absent from Mycobacterium chelonae. Antimicrob Agents Chemother 53:1367–1376.

50. Richard M, Gutiérrez AV, Kremer L. 2020. Dissecting erm(41)-Mediated Macrolide-Inducible Resistance in Mycobacterium abscessus. Antimicrob Agents Chemother 64:e01879–19.

51. Vianna JS, Machado D, Ramis IB, Silva FP, Bierhals DV, Abril MA, von Groll A, Ramos DF, Lourenço MCS, Viveiros M, da Silva PEA. 2019. The Contribution of Efflux Pumps in Mycobacterium abscessus Complex Resistance to Clarithromycin. Antibiot Basel Switz 8:153.

52. Guo Q, Chen J, Zhang S, Zou Y, Zhang Y, Huang D, Zhang Z, Li B, Chu H. 2020. Efflux Pumps Contribute to Intrinsic Clarithromycin Resistance in Clinical, Mycobacterium abscessus Isolates. Infect Drug Resist 13:447–454.

53. Ung KL, Alsarraf HMAB, Olieric V, Kremer L, Blaise M. 2019. Crystal structure of the aminoglycosides N-acetyltransferase Eis2 from Mycobacterium abscessus. FEBS J 286:4342–4355.

54. Dillon SC, Dorman CJ. 2010. Bacterial nucleoid-associated proteins, nucleoid structure and gene expression. Nat Rev Microbiol 8:185–195.

55. Kahramanoglou C, Seshasayee ASN, Prieto AI, Ibberson D, Schmidt S, Zimmermann J, Benes V, Fraser GM, Luscombe NM. 2011. Direct and indirect effects of H-NS and Fis on global gene expression control in Escherichia coli. Nucleic Acids Res 39:2073–2091.

56. Bussi C, Gutierrez MG. 2019. Mycobacterium tuberculosis infection of host cells in space and time. FEMS Microbiol Rev 43:341–361.

57. Hooppaw AJ, McGuffey JC, Di Venanzio G, Ortiz-Marquez JC, Weber BS, Lightly TJ, van Opijnen T, Scott NE, Cardona ST, Feldman MF. 2022. The Phenylacetic Acid Catabolic Pathway Regulates Antibiotic and Oxidative Stress Responses in Acinetobacter. mBio 13:e0186321.

58. Hunt TA, Kooi C, Sokol PA, Valvano MA. 2004. Identification of Burkholderia cenocepacia genes required for bacterial survival in vivo. Infect Immun 72:4010–4022.

59. Elsen S, Efthymiou G, Peteinatos P, Diallinas G, Kyritsis P, Moulis J-M. 2010. A bacteria-specific 2[4Fe-4S] ferredoxin is essential in Pseudomonas aeruginosa. BMC Microbiol 10:271.

60. Py B, Barras F. 2014. [Iron and sulfur in proteins. How does the cell build Fe-S clusters, cofactors essential for life?]. Med Sci MS 30:1110–1122.

61. Kwak N, Dalcolmo MP, Daley CL, Eather G, Gayoso R, Hasegawa N, Jhun BW, Koh W-J, Namkoong H, Park J, Thomson R, van Ingen J, Zweijpfenning SMH, Yim J-J. 2019. M ycobacterium abscessus pulmonary disease: individual patient data meta-analysis. Eur Respir J 54:1801991.

62. Bastian S, Veziris N, Roux A-L, Brossier F, Gaillard J-L, Jarlier V, Cambau E. 2011. Assessment of clarithromycin susceptibility in strains belonging to the Mycobacterium abscessus group by erm(41) and rrl sequencing. Antimicrob Agents Chemother 55:775–781.

63. Colangeli R, Helb D, Vilchèze C, Hazbón MH, Lee C-G, Safi H, Sayers B, Sardone I, Jones MB, Fleischmann RD, Peterson SN, Jacobs WR, Alland D. 2007. Transcriptional regulation of multi-drug tolerance and antibiotic-induced responses by the histone-like protein Lsr2 in M. tuberculosis. PLoS Pathog 3:e87.

64. Navarre WW, McClelland M, Libby SJ, Fang FC. 2007. Silencing of xenogeneic DNA by H-NS-facilitation of lateral gene transfer in bacteria by a defense system that recognizes foreign DNA. Genes Dev 21:1456–1471.

65. Gordon BRG, Li Y, Cote A, Weirauch MT, Ding P, Hughes TR, Navarre WW, Xia B, Liu J. 2011. Structural basis for recognition of AT-rich DNA by unrelated xenogeneic silencing proteins. Proc Natl Acad Sci U S A 108:10690–10695.

66. Summers EL, Meindl K, Usón I, Mitra AK, Radjainia M, Colangeli R, Alland D, Arcus VL. 2012. The Structure of the Oligomerization Domain of Lsr2 from Mycobacterium tuberculosis Reveals a Mechanism for Chromosome Organization and Protection. PLoS ONE 7:e38542.

67. Medjahed H, Singh AK. 2010. Genetic manipulation of Mycobacterium abscessus. Curr Protoc Microbiol Chapter 10:Unit 10D.2.

68. Chen S. 2023. Ultrafast one-pass FASTQ data preprocessing, quality control, and deduplication using fastp. iMeta 2:e107.

69. Dobin A, Davis CA, Schlesinger F, Drenkow J, Zaleski C, Jha S, Batut P, Chaisson M, Gingeras TR. 2013. STAR: ultrafast universal RNA-seq aligner. Bioinformatics 29:15– 21.

70. Liao Y, Smyth GK, Shi W. 2014. featureCounts: an efficient general purpose program for assigning sequence reads to genomic features. Bioinformatics 30:923–930.

71. Varet H, Brillet-Guéguen L, Coppée J-Y, Dillies M-A. 2016. SARTools: A DESeq2- and EdgeR-Based R Pipeline for Comprehensive Differential Analysis of RNA-Seq Data. PloS One 11:e0157022.

72. Langmead B, Salzberg SL. 2012. Fast gapped-read alignment with Bowtie 2. 4. Nat Methods 9:357–359.

73. Danecek P, Bonfield JK, Liddle J, Marshall J, Ohan V, Pollard MO, Whitwham A, Keane T, McCarthy SA, Davies RM, Li H. 2021. Twelve years of SAMtools and BCFtools. GigaScience 10:giab008.

74. Robinson JT, Thorvaldsdóttir H, Winckler W, Guttman M, Lander ES, Getz G, Mesirov JP. 2011. Integrative genomics viewer. Nat Biotechnol 29:24–26.

75. Ramírez F, Ryan DP, Grüning B, Bhardwaj V, Kilpert F, Richter AS, Heyne S, Dündar F, Manke T. 2016. deepTools2: a next generation web server for deep-sequencing data analysis. Nucleic Acids Res 44:W160–5.

76. Zhang Y, Liu T, Meyer CA, Eeckhoute J, Johnson DS, Bernstein BE, Nusbaum C, Myers RM, Brown M, Li W, Liu XS. 2008. Model-based analysis of ChIP-Seq (MACS). Genome Biol 9:R137.

77. Quinlan AR, Hall IM. 2010. BEDTools: a flexible suite of utilities for comparing genomic features. Bioinforma Oxf Engl 26:841–842.

78. Taboada B, Estrada K, Ciria R, Merino E. 2018. Operon-mapper: a web server for precise operon identification in bacterial and archaeal genomes. Bioinforma Oxf Engl 34:4118–4120.

79. Stark R, Brown G. 2011. DiffBind: differential binding analysis of ChIP-Seq peak data. R Package Version 100.

80. Woods GL, Brown-Elliott BA, Conville PS, Desmond EP, Hall GS, Lin G, Pfyffer GE, Ridderhof JC, Siddiqi SH, Wallace RJ, Warren NG, Witebsky FG. 2011. Susceptibility Testing of Mycobacteria, Nocardiae, and Other Aerobic Actinomycetes, 2nd ed. Clinical and Laboratory Standards Institute, Wayne (PA). http://www.ncbi.nlm.nih.gov/books/NBK544374/. Retrieved 9 November 2022.

81. Halloum I, Viljoen A, Khanna V, Craig D, Bouchier C, Brosch R, Coxon G, Kremer L. 2017. Resistance to Thiacetazone Derivatives Active against Mycobacterium abscessus Involves Mutations in the MmpL5 Transcriptional Repressor MAB_4384. Antimicrob Agents Chemother 61:e02509–16.

82. Le Run E, Arthur M, Mainardi J-L. 2018. In Vitro and Intracellular Activity of Imipenem Combined with Rifabutin and Avibactam against Mycobacterium abscessus. Antimicrob Agents Chemother 62:e00623–18.

83. Bakala N’Goma JC, Le Moigne V, Soismier N, Laencina L, Le Chevalier F, Roux A- L, Poncin I, Serveau-Avesque C, Rottman M, Gaillard J-L, Etienne G, Brosch R, Herrmann J-L, Canaan S, Girard-Misguich F. 2015. Mycobacterium abscessus phospholipase C expression is induced during coculture within amoebae and enhances M. abscessus virulence in mice. Infect Immun 83:780–791.

84. Le Moigne V, Blouquit-Laye S, Desquesnes A, Girard-Misguich F, Herrmann J-L. 2022. Liposomal amikacin and Mycobacterium abscessus: intimate interactions inside eukaryotic cells. J Antimicrob Chemother dkac348.

