## Supplemental figure S1, S2, S3; Supplemental table S1, S2, S3 for "Lsr2, a pleiotropic regulator at the core of the infectious strategy of *Mycobacterium abscessus*"

**Supplementary data:**

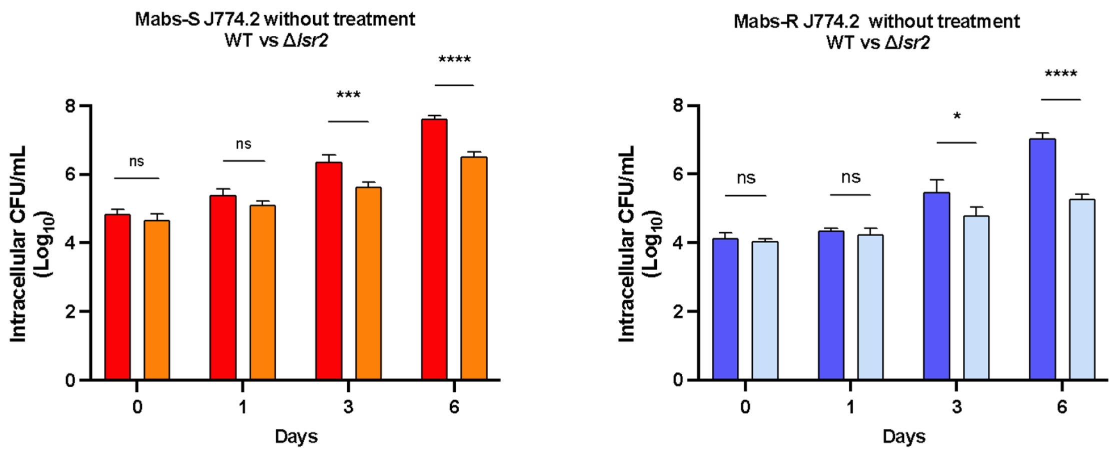


**Figure S1: Intracellular growth of *M. abscessus* Mabs-S and Mabs-R *wild-type and lsr2* mutant strains in macrophages as control without antibiotics treatment.** Murine J774.2 macrophages were infected with Mabs-S-WT (red), Mabs-S-Δ*lsr2* (orange), Mabs-R-WT (blue) and Mabs-R-Δ*lsr2* (light blue) at an MOI of 10. Intracellular growth was evaluated by counting CFUs at various time points post-infection (days 0, 1, 3, and 6). Data are representative of three independent experiments and represent means ± SEM. Differences between means were analyzed by two-way ANOVA and the Tukey post-test, allowing multiple comparisons. ns, non-significant, **P < 0.05, ***P < 0.001, and ****P < 0.0001.*

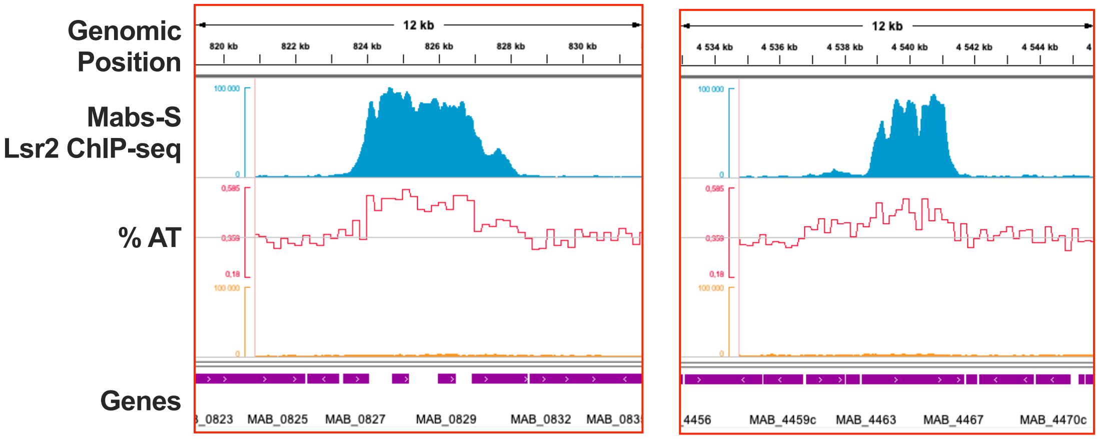


**Figure S2: Lsr2 binds specifically on AT-rich sequences of the M. abscessus genome.** Distribution of Lsr2 (in blue) is strongly correlated with enhanced AT content as exemplified for genomic regions encompassing genes from *MAB_0827* to *MAB_0842* and from *MAB_4463* to *MAB_4467*. Raw sequencing coverage from Input sample is also pictured in orange as a control for Lsr2 binding specificity. The bottom part corresponds to gene positions.

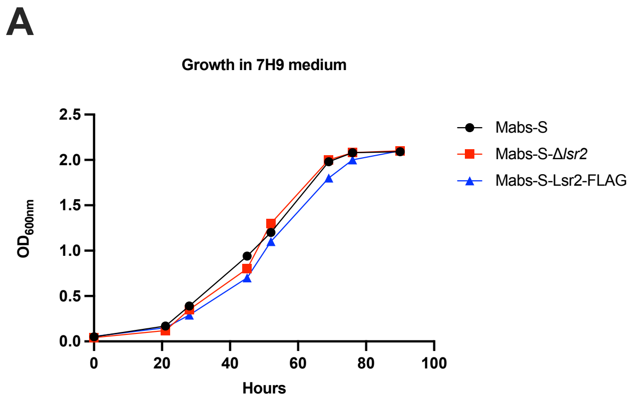


**Figure S3: (A)** Growth curve showing the absence of a growth defect in Mabs-S-Δ*lsr2* and Mabs-S-Lsr2-FLAG strains as compared to Mabs-S. **(B)** Growth curve showing the absence of a growth defect in Mabs-R-Δ*lsr2* and Mabs-R-Lsr2-FLAG strains as compared to Mabs-R.

**Table S1. Primers used in this study**

| Name | Sequence |
|---|---|
| *5'_Homology_Lsr2_region_Fwd* | 5'-aatgtatccgtcgttgtcagaag-3' |
| *5'_Homology_Lsr2_region_Rev* | 5'-tcctttttccttccaaaagcg-3' |
| *Lsr2-3XFLAG-zeocin_Fwd* | 5'- atggctaagaaggtcacgg-3' |
| *Lsr2-3XFLAG-zeocin_Rev* | 5'-ggtctgacgctcagtggaac-3' |
| *3'_Homology_Lsr2_region_Fwd* | 5'-tacgttcccgggacaatttc-3' |
| *3'_Homology_Lsr2_region_Rev* | 5'-ctcgaaatcaccgcggtag-3' |
| *SigA_Fwd* | 5'-tccgagaaagacaaggcttc-3' |
| *SigA_Rev* | 5'-ccagctcgacttcctcttcg-3' |
| *MAB_2355c_Fwd* | 5'-atgccaaactcgcgaacta-3' |
| *MAB_2355c_Rev* | 5'-tctgccggtacatcaacacc-3' |
| *erm41_Fwd* | 5'-ggaagatgtccggatagcgg-3' |
| *erm41_Rev* | 5'-gtatcagtgcgctggtgact-3' |
| *eis2_Fwd* | 5'-gtgttgtgagcgatgcgatc-3' |
| *eis2_Rev* | 5'-cggaagaaagtcgtcggtca-3' |
| *MAB_1409c_Fwd* | 5'-gtcgatcttctccgacgtcc-3' |
| *MAB_1409c_Rev* | 5'-gtcatcgcaaggatcgggat-3' |
| *lsr2_Fwd* | 5'-gagaccgttgaattcggtg-3' |
| *lsr2_Rev* | 5'-gctgattacgcagcttctcc-3' |
| *MmpL8_Fwd* | 5'-ctcgaatcagaccctgacgttcat-3' |
| *MmpL8_Rev* | 5'-tgcccaacttggtgaatcccat-3' |
| *MAB_2037_Fwd* | 5'-gcagacacgcatggcattaagt-3' |
| *MAB_2037_Rev* | 5'-gaaaccgtagagcgacctgaaagt-3' |

**Table S2. Differentially expressed genes between Mabs-S and Mabs-R morphotypes of *M. abscessus***

| Genes | Mab-S vs Mabs-R | | Functions of encoded proteins |
|---|---|---|---|
| | $Log_2FC$ | Adj. p-value | |
| ***mps1*** | -1.724 | 3.06E-03 | Glycopeptidolipid biosynthesis protein |
| ***mps2*** | -3.881 | 6.15E-15 | Glycopeptidolipid biosynthesis protein |
| ***gap*** | -2.633 | 4.79E-06 | Integral membrane protein |
| ***MAB_2552c*** | -1.546 | 3.51E-15 | Lipid transport and metabolism protein |
| ***MAB_4272c*** | 2.477 | 0.005783288 | Molecular chaperone GrpE (HSP-70 cofactor) protein |
| ***MAB_4273c*** | 2.462 | 0.004734461 | Molecular chaperone DnaK (HSP-70) protein |
| ***MAB_1242c*** | 2.121 | 0.000254187 | Hypothetical protein |
| ***MAB_1243c*** | 1.875 | 0.005783288 | Hypothetical protein |
| ***MAB_1247c*** | 2.381 | 0.026330185 | Hypothetical protein |
| ***nrdF*** | 2.031 | 0.026330185 | Nucleotide transport and metabolism protein |
| ***nrdE*** | 1.894 | 0.005783288 | Nucleotide transport and metabolism |
| ***nrdI*** | 1.851 | 0.026330185 | Nucleotide transport and metabolism |
| ***nrdH*** | 1.677 | 0.034602339 | Posttranslational modification protein |

**Table S3. Regulation of *mmpL8*$_{MAB}$ locus by Lsr2 in Mabs-S and Mabs-R morphotypes of *M. abscessus***

| *mmpL8*$_{MAB}$ locus (genes) | Mabs-S *Δlsr2* vs WT | | Mabs-S *Δlsr2* vs WT | |
|---|---|---|---|---|
| | $Log_2FC$ | Adj. p-value | $Log_2FC$ | Adj. p-value |
| ***papA2*** | -6.362 | 6.08E-21 | 2.117 | 1.46E-02 |
| ***lipP*** | -6.412 | 1.07E-23 | 2.046 | 1.15E-02 |
| ***MAB_0854*** | -5.868 | 3.29E-18 | 1.949 | 2.60E-02 |
| ***mmpL8*** | -5.886 | 7.64E-17 | 2.174 | 1.89E-02 |
| ***tetR*** | -6.002 | 1.04E-06 | 4.656 | 2.27E-36 |
| ***MAB_0857*** | -5.381 | 4.42E-20 | 4.342 | 2.25E-34 |
| ***MAB_0858*** | -6.21 | 6.15E-50 | 1.323 | 5.60E-03 |
| ***MAB_0859*** | -5.393 | 1.83E-27 | 1.041 | 1.76E-05 |
